# Adaptive shifts in gene regulation underlie a developmental delay in thermogenesis in high-altitude deer mice

**DOI:** 10.1101/2019.12.17.880112

**Authors:** Jonathan P. Velotta, Cayleih E. Robertson, Rena M. Schweizer, Grant B. McClelland, Zachary A. Cheviron

## Abstract

Aerobic performance is tied to fitness as it influences an animal’s ability to find food, escape predators, or survive extreme conditions. At high altitude, where low O_2_ availability and persistent cold prevail, maximum metabolic heat production (thermogenesis) is an aerobic performance trait that is intimately linked to survival. Understanding how thermogenesis evolves to enhance survival at high altitude will yield insight into the links between physiology, performance, and fitness. Recent work in deer mice (*Peromyscus maniculatus*) has shown that adult mice native to high-altitude have higher thermogenic capacities under hypoxia compared to lowland conspecifics, but developing high-altitude pups delay the onset of thermogenesis. This suggests that natural selection on thermogenic capacity varies across life stages. To determine the mechanistic cause of this ontogenetic delay, we analyzed the transcriptomes of thermo-effector organs – brown adipose tissue and skeletal muscle – in developing deer mice native to low- and high-altitude. We demonstrate that the developmental delay in thermogenesis is associated with adaptive shifts in the expression of genes involved in nervous system development, fuel/O_2_ supply, and oxidative metabolism gene pathways. Our results demonstrate that selection has modified the developmental trajectory of the thermoregulatory system at high altitude and has done so by acting on the regulatory systems that control the maturation of thermo-effector tissues. We suggest that the cold and hypoxic conditions of high altitude may force a resource allocation trade-off, whereby limited energy is allocated to developmental processes such as growth, versus active thermogenesis during early development.

## Introduction

Fitness in the wild is determined by suites of interacting traits that influence variation in whole-organism performance. This is because performance determines an organism’s ability to conduct ecologically-relevant tasks such as avoiding predators, competing for resources, and surviving extreme events (Huey and Stevenson 1979; Garland and Losos 1994; Irschick and Garland 2001; Irschick et al. 2008; Campbell-Staton et al. 2017). Many of these ecologically-relevant tasks that impinge on whole-organism performance are ultimately dependent upon the capacity for aerobic metabolism, and thus, the ability to transport and utilize O_2_. As such, the development of the physiological machinery needed to delivery O_2_ and metabolize fuels is intimately tied to both postnatal juvenile survival and the future reproductive success as adults. The timing of key developmental events, and responses to early environmental exposures, should therefore influence the evolution of whole-organism performance (Lailvaux and Husak 2014). Studies that seek to understand how performance evolves in the context of development are rare but are needed to determine the causal connections between phenotype and fitness in the wild.

Extreme environments, such as high-altitude habitats > 3,000 meters above sea level (a.s.l. Bouverot 1985), are windows into the mechanisms that shape adaptive variation in performance, since the selection pressures are few in number and strong in magnitude (Garland and Carter 1994). The agents of selection at high altitude – cold temperature and unavoidable reductions in available O_2_ (hypobaric hypoxia) – have led to clear examples of local adaptation in animals that live there permanently (e.g., Beall 2007; Storz et al. 2010; Simonson et al. 2011). A considerable amount of recent work on high-altitude adaptation has been conducted on the North American deer mouse (*Peromyscus maniculatus*) owing to its broad altitudinal distribution, ranging from sea level to >4,000 m a.s.l. (King 1968). Recent comparative physiological work demonstrates that adult high-altitude deer mice from the Colorado Rocky Mountains have consistently higher aerobic performance capabilities than conspecifics in the Great Plains: high-altitude deer mice are capable of greater aerobic performance under hypoxia and in response to extreme cold (referred to as thermogenic capacity) measured in the wild and in the laboratory (Cheviron et al. 2012; Cheviron et al. 2013; Cheviron et al. 2014; Lui et al. 2015; Lau et al. 2017; Tate et al. 2017). Capture-mark-recapture data confirm that higher thermogenic capacities are beneficial for survival at high-altitude (Hayes and O’Connor 1999). Available data suggests that enhanced thermogenic capacity provides an important performance benefit in high-altitude deer mice that is strongly tied to fitness during adulthood.

Recent work has uncovered the factors that shape adaptive variation in thermogenic performance to illuminate links between physiology and fitness. This work, conducted in adult deer mice, suggests that improved thermogenic performance is related to alterations to O_2_ transport and utilization, including more effective breathing patterns (Ivy and Scott 2017), higher blood-O_2_ affinity (Snyder 1981; Snyder et al. 1982; Chappell and Snyder 1984; Storz 2007; Storz 2016) and circulation (Tate et al. 2017), an improved capacity to oxidize lipids as fuel (Cheviron et al. 2012; Cheviron et al. 2014), and a greater oxidative capacity in skeletal muscles and their mitochondria (Lui et al. 2015; Scott et al. 2015; Mahalingam et al. 2017). Despite the importance of these changes in adults, it is not clear how high-altitude animals have evolved to survive the unique challenges of cold and hypoxic stress during development. Early post-natal development, however, is critically linked to fitness: altricial rodent pups are small, immobile, and reliant on limited energy supplied through maternal care, and pup mortality can be extremely high in the wild (Hill 1983). Moreover, mouse pups are not born with the ability to generate heat (Pembrey 1895). Independent thermogenic abilities develop as a result of the maturation of thermo-effector organs, brown adipose tissue (BAT) and skeletal muscle, which permits non-shivering and shivering thermogenesis, respectively (Lagerspetz 1966; Arjamaa and Lagerspetz 1979). It follows than that if selection pressures on thermogenic performance are constant across life stages, then thermogenesis should develop faster in high-altitude pups to support an improved adult performance.

Robertson et al. (2019) however have recently demonstrated that non-shivering thermogenesis (NST) is delayed by approximately 2-days in high-altitude deer mouse pups compared to lowland conspecifics and a closely related, but strictly lowland species, *P. leucopus*. This delay in the onset of NST is also further associated with a delay in the onset of shivering thermogenesis until the normal date of weening (Robertson and McClelland 2019; Fig. 1). The authors suggest that the observed ontogenetic delay may be an evolutionary response to limitations in energy production or allocation at high altitude (*e.g.*, as a result of low O_2_), which force a trade-off between active thermogenesis vs. other developmental functions such as growth. Consistent with this interpretation, growth rates under common conditions are identical between low- and high-altitude pups, despite the fact that high-altitude mothers produce larger litters (Robertson et al. 2019).

**Figure 1.**
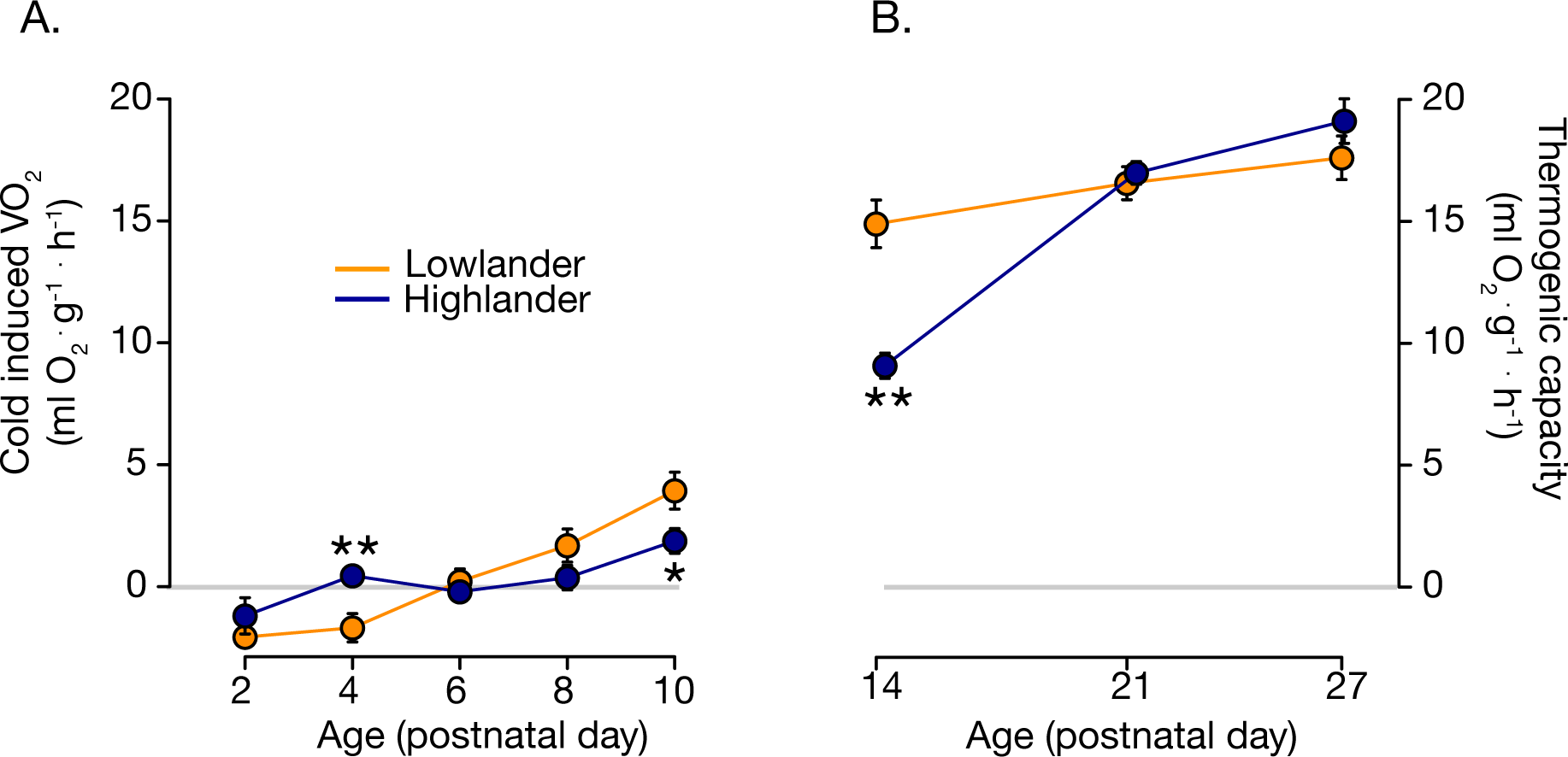
The development of thermogenesis is delayed in highland (blue) related to lowland (orange) deer mice. (A) Cold (24°C)-induced O_2_ consumption rate (VO_2_) across postnatal day p2-p10. Values constitute the difference between cold-induced and normothermic VO_2_, which controls for the stress of separating pups from their mothers at a young age. Values crossing zero indicates that pups are able to generate heat independently. (B) Thermogenic capacity (VO_2_ max during exposure to 5°C in heliox) under normoxia is significantly lower in highlanders than lowlanders at p14. Values represent mean mass-corrected VO_2_. *p<0.05; **p<0.01. Data from Robertson et al. 2019 (A) and Robertson and McClelland 2019 (B).

We tested the hypothesis that the ontogenetic delay in thermogenesis is driven by regulatory changes that delay the development of the primary thermo-effector organs. To do this, we compared BAT and skeletal muscle transcriptomes in deer mice native to low and high altitudes across the first 27 days of life. We associated variation in transcript abundances to variation in thermogenic capabilities using a gene co-expression network approach that has been used to link gene regulation to physiological function in variety of evolutionary contexts (Whitehead 2012; Cheviron et al. 2014; DeBiasse and Kelly 2016; Velotta et al. 2017; Campbell-Staton et al. 2018). We show that the delay in thermogenesis in high-altitude mice is associated with a delay in the expression of gene networks that function in nervous system control of, and fuel/O_2_ supply to, BAT, as well as aerobic metabolism and mitochondrial biogenesis in skeletal muscle. Finally, using a combination of phenotypic divergence and population-genetic approaches, we provide evidence that the ontogenetic delays in thermogenesis and their associated regulatory changes are adaptive at high altitude. By combining these approaches, we demonstrate, for the first time, that selection has altered the developmental trajectories of a thermoregulatory system by acting on the regulatory control of thermo-effector organ phenotype.

## Results

### The ontogeny of thermogenesis

We reanalyzed cold-induced metabolic rate data (rates of O_2_ consumption, VO_2_) from Robertson et al. (2019) and Robertson and McClelland (2019) to assess the ontogeny of thermogenesis in deer mice native to lowland (320 m a.s.l) and highland (4,350 m a.s.l) habitats. VO_2_ of pups post-natal age p0-p10 was measured during exposure to a mild cold stressor (24°C), and was calculated as the difference in cold-induced VO_2_ compared to that of normothermic (30°C) littermates (see Materials and Methods). VO_2_ in these young pups was ~0 until p6 for highlanders and lowlanders (Fig. 1A). After p6, cold-induced VO_2_ increased in both populations, but did so at a slower rate in highlanders (Fig. 1). Statistically significant population differences in VO_2_ were detected at p4 and p10 (Fig. 1). We measured thermogenic capacity in older pups (age p14, p21 and p27) as VO_2_-max during exposure to cold (4°C) in a heliox air mixture. Highland deer mice exhibited a significantly reduced thermogenic capacity at p14 compared to lowlanders (~6 ml O_2_ per gram per hour, on average), whereas by p21 thermogenic capacities between populations were nearly equivalent (Fig. 1B). Although there is a trend towards highlanders surpassing lowlanders by p27, this result was not significant (*p*=0.2).

### Transcriptional correlates of thermogenesis

We sequenced the transcriptomes of thermo-effector organs, brown adipose tissue (BAT) and skeletal muscle, in order to assess regulatory mechanisms underlying evolutionary changes in the ontogeny of thermogenesis at high-altitude. Principal components analysis (PCA) of transcriptome-wide patterns of gene expression revealed separation between lowlanders and highlanders, and across ontogeny, in BAT and skeletal muscle (Fig. 2). For BAT, highlanders and lowlanders were distinguishable along PC1, which explained 12% of total variation in expression. By contrast, PC2, which explained 10% of the variance, distinguished post-natal age (Fig. 2A). We observed similar separations in muscle samples, although in this case populations were distinguishable along PC1 (20% variance), and post-natal age distinguishable along PC2 (10% of variance). 95% confidence ellipses were drawn around each population demonstrating that populations were entirely (skeletal muscle, Fig 2B) or almost entirely (BAT Fig. 2A) non-overlapping in PC1 and PC2 space.

**Figure 2.**
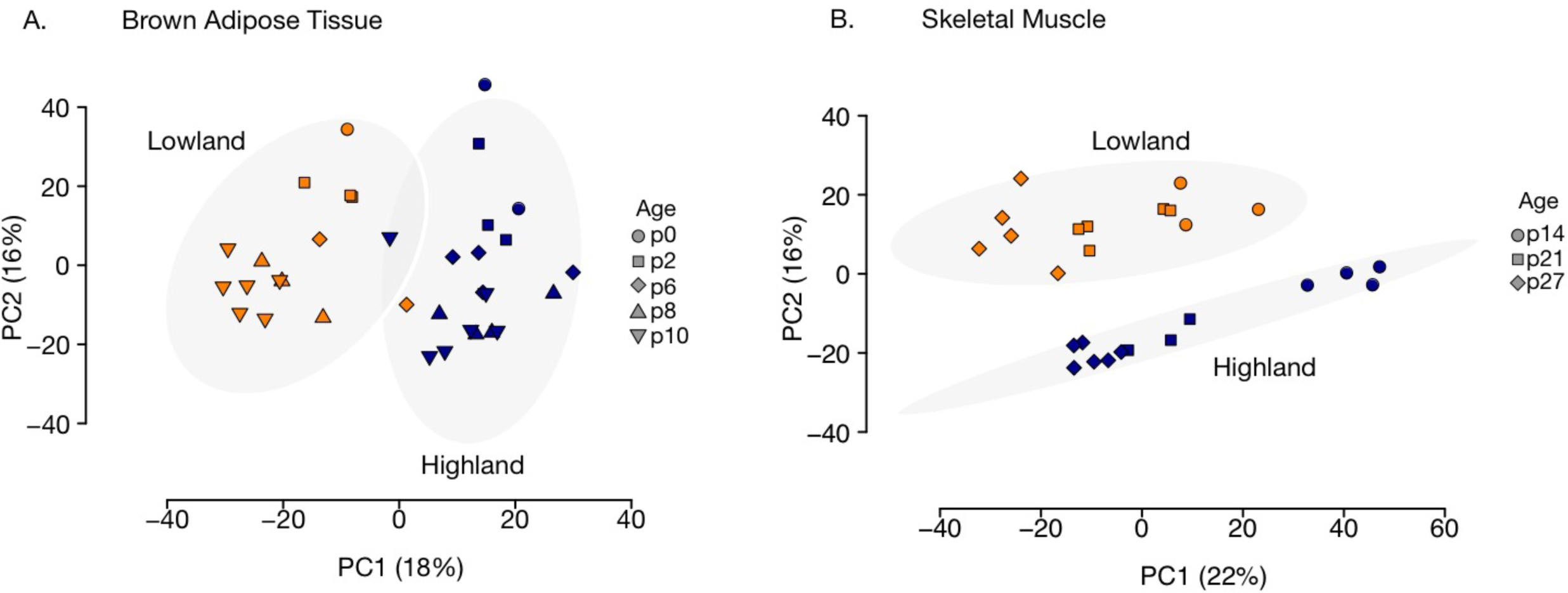
Principal components analysis (PCA) transcriptome expression data for thermo-effector organs brown adipose tissue and (A) skeletal muscle (B) in lowland (orange points) and highland (blue points) deer mice. Symbols represents ages from postnatal day p0-p10 for brown adipose tissue, and p14-p27 in skeletal muscle. Populations separated along PC1 in brown adipose tissue data and PC2 in skeletal muscle data. Grey circles are 95% confidence ellipses drawn around each population to highlight separation.

Weighted gene co-expression network analysis (WGCNA) of BAT samples across p0-p10 yielded 38 modules ranging in size from 61 to 1,073 genes (See Supplemental Table S2 for a full list of module assignments). A total of 1,588 out of 11,192 genes could not be assigned to any module (Module B0). We found ten modules exhibiting statistically significant associations between module eigengene and cold-induced VO_2_, after correction for multiple testing (Table 1). Of the ten VO_2_-associated modules, six exhibited significant population effects on module eigengene (Table 1). Two of these (modules B11 and B35) were not significantly enriched for GO terms or were enriched for terms unrelated to thermogenesis (e.g., “drug metabolism”; see Table S4 for the full list of enriched terms). Modules B4 and B12 (Table 1) were both significantly enriched for genes that encode ribosomal proteins (Supplemental Table S4), which may indicate differences in protein translation between populations. No significant interaction effects were detected for any VO_2_-associated module (Table 1).

**Table 1.** Brown adipose tissue modules significantly correlated with cold-induced VO_2_. Full association test results are presented in Supplementary Table S3. Effect of population (highland vs. lowland), age (postnatal day p0-p10), or their interaction from ANOVA models on rank-transformed module expression values (module eigengene values). Final two columns present results of Pearson correlation between module expression and cold-induced VO_2_. *P*-values from association tests and ANOVAs were corrected for multiple testing using the false-discovery rate method.

| Module | Module size | Population |  | Age |  | Pop x Age |  | VO<sub>2</sub> correlation |  |
| --- | --- | --- | --- | --- | --- | --- | --- | --- | --- |
|  |  | F<sub>1,30</sub> | *P* | F<sub>1,30</sub> | *P* | F<sub>1,30</sub> | *P* | *r* | *P* |
| B2 | 1022 | 7.4 | 0.41 | 3.2 | 1.0 | 5.36 | 1.0 | -0.49 | **0.02** |
| B3 | 773 | 27.0 | **<0.001** | 20.8 | **<0.001** | 1.67 | 1.0 | 0.67 | **<0.001** |
| B4 | 252 | 18.6 | **0.01** | 38.1 | **<0.001** | 0.43 | 1.0 | -0.56 | **0.004** |
| B11 | 99 | 162.1 | **<0.001** | 26.8 | **<0.001** | 0.04 | 1.0 | -0.47 | **0.02** |
| B12 | 605 | 21.5 | **<0.001** | 128.2 | **<0.001** | 4.90 | 1.0 | -0.81 | **<0.001** |
| B16 | 149 | 2.7 | 1.0 | 25.9 | **<0.001** | 0.06 | 1.0 | -0.55 | **0.01** |
| B32 | 267 | 0.1 | 1.0 | 32.9 | **<0.001** | 0.36 | 1.0 | -0.52 | **0.01** |
| B33 | 1073 | 0.7 | 1.0 | 53.9 | **<0.001** | 0.85 | 1.0 | 0.64 | **<0.001** |
| B35 | 74 | 37.3 | **<0.001** | 0.9 | 1.0 | 0.02 | 1.0 | -0.45 | **0.03** |
| B36 | 708 | 26.5 | **<0.001** | 51.5 | **<0.001** | 1.52 | 1.0 | 0.58 | **<0.001** |

Modules B3 and B36, by contrast, were positively associated with cold-induced VO_2_ (Fig. 3A-B), expressed at significantly lower levels in highlanders across development (Fig. 3B-C), and exhibited significant enrichment for functions that reflect a delay in the development of BAT in highlanders (Fig. 3E-F). Module B3 was enriched for functional terms related to fatty acid metabolism in the mitochondria, including the GO Biological Process term “fatty acid metabolic process”, the GO Cellular Component term “mitochondrion,” the KEGG pathway “fatty acid degradation (Fig. 3E), and Reactome and WikiPathways terms “fatty acid metabolism” and “fatty acid beta oxidation (Supplementary Table S4).” Module B36 was enriched for terms related to the vascularization of BAT and the development of its neural circuitry (Fig. 3F): vascularization terms include the GO Biological Processes “blood vessel development” and “vasculature development,” as well as the KEGG pathway “VEGF signaling,” which is the major pathway regulating new blood vessel growth. Indeed, *Vegfa* exhibits significantly lowered expression in highland compared to lowland mice (Fig. 3F inset), though it was not assigned to module B36 (Supplemental Table S2). Finally, we detected enrichment of the KEGG pathway “axon guidance,” which represents a key stage in the development of neuronal networks. Fig. 3B-C reveal that expression values for both of these modules increased from birth to post-natal day 10, and that expression was consistently higher among lowlanders at nearly every time point (*p*<0.001 population effect for both modules).

**Figure 3.**
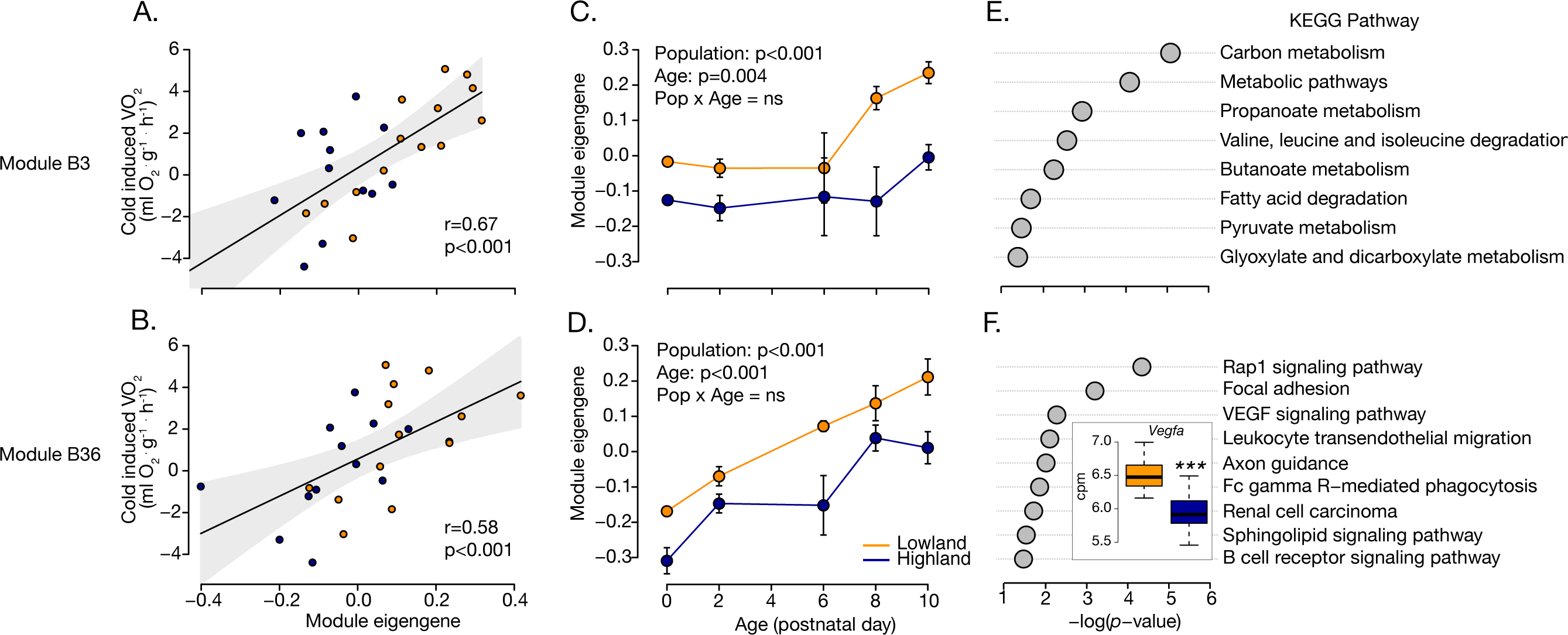
Two brown adipose tissue modules are associated with variation in thermogenesis in p0-p10 deer mouse pups. (A-B): panels show the significant positive association between module eigengene and cold-induced VO_2_ (measure of thermogenesis). Eigengene values of modules B3 (C) and B36 (D) are significantly affected by age, and significantly differentiated between lowland (orange) and highland (blue) populations. We detected significant enrichment of KEGG pathways related to metabolism and fatty acid degradation in B3 (E), and VEGF signaling and axon guidance in B36, among others (F). Values in (E-F) are the negative log of the enrichment test *p*-value after correction for multiple testing using the g:SCS algorithm in *gProfilerR* (Reimend et al. 2016). Inset in (F) shows average expression (log counts per million; cpm) across age for *Vegfa* in lowlanders (orange) vs. highlanders (blue). *** p<0.001. See Table S3 for full functional enrichment analysis results.

We performed WGCNA on skeletal muscle transcriptomes from lowland and highland deer mice sampled at postnatal days 14, 21, and 27. WGCNA identified seven modules ranging in size from 92 to 2,878 genes (Table 2). A total of 3,529 out of 11,104 genes could not be assigned to any module (M0; Supplemental Table S2). Although individual-level thermogenic capacity data do not exist for the samples sequenced, we did detect significant correlations between module eigengene and thermogenic capacity at the level of population and age for two modules, M2 (r=−0.94; *p*=0.006) and M6 (r=0.98; *p*<0.001; Fig. 4A; Table 2). Although we detected an overall effect of postnatal age, but not population, on both of these modules (Table 2), expression followed a pattern that closely resembled population-level variation in thermogenic capacity. For module M6 in particular, as with thermogenic capacity (Fig 1B), expression was lower in highland compared to lowland deer mice at p14 (F_1,4_ = 27.7; *p*=0.006), but no different from lowlanders at p21 or p27 (*p*>0.05; Fig. 4B). The opposite was true of M2, whereby module expression at p14 was significantly higher in high-altitude mice (F_1,4_ = 27.7; *p*=0.02).

**Table 2.** Association-test and ANOVA results on skeletal muscle modules. Effect of population (highland vs. lowland), ontogenetic age (postnatal day p14-p27), or their interaction on rank-transformed module expression (module eigengene values) is presented. Final two columns present results of Pearson correlation between module expression and thermogenic capacity. Note that correlations were performed on population-age means since VO_2_ measurements did not exist for the individuals sequenced. *P*-values from association tests and ANOVAs were corrected for multiple testing using the false-discovery rate method.

| Module | Module size | Population |  | Age |  | Pop x Age |  | VO<sub>2</sub> correlation |  |
| --- | --- | --- | --- | --- | --- | --- | --- | --- | --- |
|  |  | F<sub>1,22</sub> | *P* | F<sub>1,22</sub> | *P* | F<sub>1,22</sub> | *P* | *r* | *P* |
| M1 | 92 | 3.4 | 0.6 | 15.0 | 0.006 | 2.4 | 1 | 0.18 | 0.99 |
| M2 | 2654 | 1.0 | 1 | 150.5 | <0.001 | 0.3 | 1 | -0.94 | **0.02** |
| M3 | 1052 | 11.9 | 0.02 | 1.8 | 1 | 2.3 | 1 | -0.79 | 0.15 |
| M4 | 288 | 97.8 | <0.001 | 8.7 | 0.06 | 1.7 | 1 | 0.0002 | 0.99 |
| M5 | 168 | 4.6 | 0.4 | 3.5 | 0.61 | 4.4 | 0.4 | 0.19 | 0.99 |
| M6 | 2878 | 3.2 | 0.7 | 64.4 | <0.001 | 0.1 | 1 | 0.98 | **0.004** |
| M7 | 443 | 91.1 | <0.001 | 7.2 | 0.11 | 1.0 | 1 | 0.07 | 0.99 |

**Figure 4.**
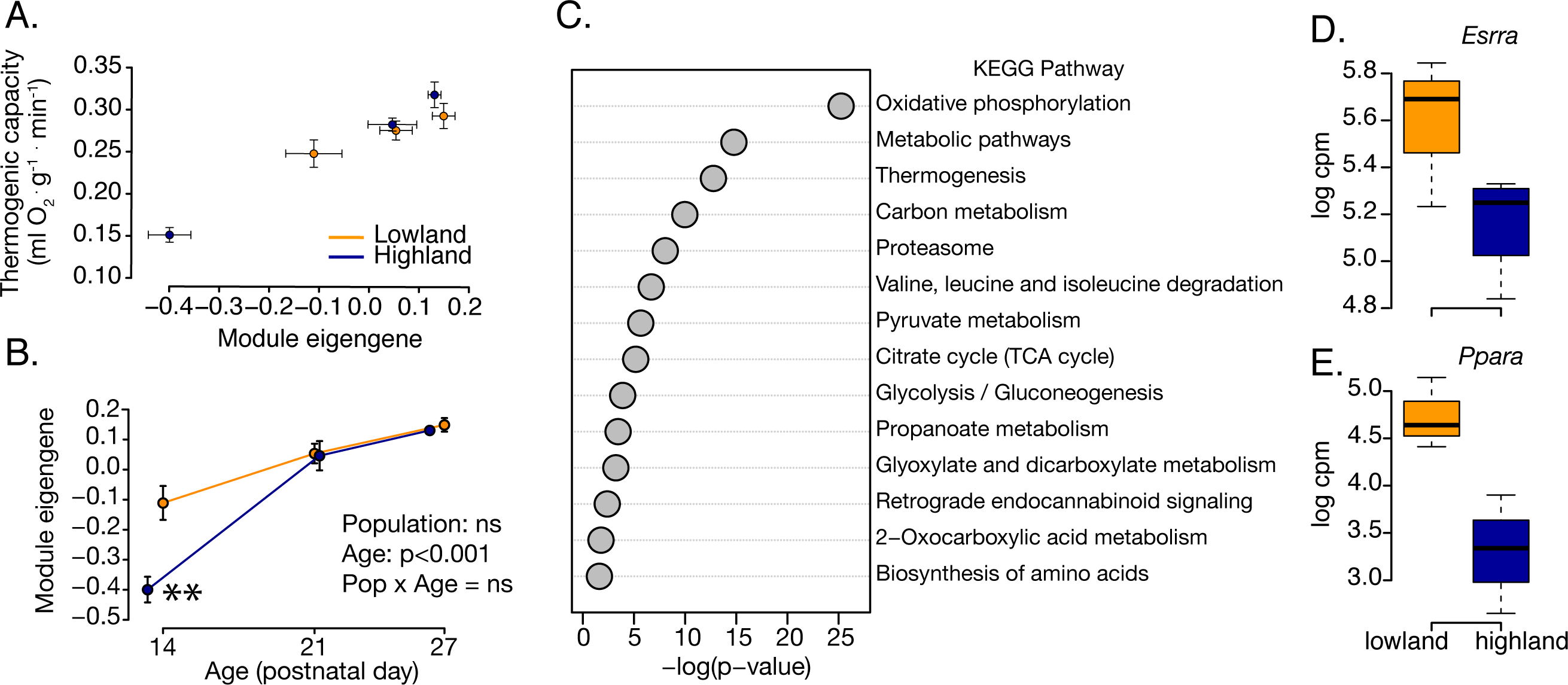
Skeletal muscle module M6 is significantly associated with thermogenic capacity in lowland (orange) and highland (blue) deer mice aged p14-p27. (A) Module expression is positively correlated with thermogenic capacity at the population/age level (individual level data not available). (B) Highland deer mice exhibit significantly lower module eigengene values at postnatal day p14 (** p<0.01), and no differences in expression at p21 or p27. (C) Functional enrichment analysis reveals enrichment of KEGG pathways involved in aerobic metabolism and thermogenesis among others. Values are the negative log of the enrichment test *p*-value after correction for multiple testing using the g:SCS algorithm in *gProfilerR* (Reimend et al. 2016). Expression values (log counts per million; cpm) for the transcription factors *Esrra* (D) and *Ppara* (E) at p14 are lower for highlanders compared to lowlanders; both transcription factors are involved in the regulation of mitochondrial biogenesis and metabolic pathways.

Functional enrichment analysis of VO_2_-associated skeletal muscle modules revealed enrichment of a wide variety of biological functions (Supplemental Table S4). Module M2 for example was enriched for pathways that function in the development of muscle blood supply, including the GO Biological Process terms “angiogenesis, blood vessel morphogenesis, and vasculogenesis.” This result may reflect an initiation of blood vessel development that eventually leads to a greater production of capillaries in highlanders by p21 (Robertson and McClelland 2019). Indeed, along with a greater proportion of oxidative fiber types, increased capillary density is thought to be an adaptive advantage at high-altitude that contributes to improved aerobic performance capabilities in adult mice (Lui et al. 2015; Scott et al. 2015; Nikel et al. 2018).

By contrast to M2, module M6 was enriched for functions related to energy metabolism and the generation of ATP, and thermogenesis, including the KEGG terms “oxidative phosphorylation, thermogenesis, citrate cycle, and glycolysis/gluconeogenesis (Fig. 4C).” Enrichment of the GO Cellular Component terms “NADH dehydrogenase complex and mitochondrial respiratory chain complex I” indicates that energy metabolism functions are related to changes in the production of NADH:ubiquinone oxidoreductase/mitochondrial complex I, the first protein complex in the electron transport chain. We also detected significant enrichment of transcription factor binding sites for estrogen-related receptor alpha, ERR1 (Table S4; *Esrra* in the *Peromyscus* genome), which is a major regulator of mitochondrial biogenesis, gluconeogenesis, oxidative phosphorylation, and gluconeogenesis. Indeed, the *Esrra* gene itself was assigned to module M6 and its expression at p14 is strongly downregulated in highlanders relative to lowlanders (Fig. 4D). Detection of *Ppara* (peroxisome proliferator activated receptor alpha) in module M6 (Fig. 4E) suggests that fatty acid oxidation may also be depressed in highlanders, since this gene is a major regulator of fat metabolism in muscle (Issemann and Green 1990).

### Evidence of selection on candidate transcriptional modules

Among the transcriptional modules identified by WGCNA, three (B3, B36, M6) were significantly and positively associated with variation in VO_2_. These modules thus represent candidate pathways underlying the delay in thermogenesis in high-altitude mice. We took a two-pronged approach to determining whether developmental delays in candidate module expression are adaptive at high-altitude. First, we calculated P_ST_ (an estimate of phenotypic differentiation following Brommer 2011) on module eigengene values of candidate modules to test whether divergence between lowland and highland populations exceeded neutral expectations as set by F_ST_ (an estimate of neutral genetic differentiation). For BAT modules B3 and B36, P_ST_ exceeded F_ST_ at nearly all values of *c/h*^*2*^ (Fig. 5A-B); *c/h*^*2*^ represents the unknown degree to which phenotypes are due to additive genetic effects. Brommer (2011) assumes a null *c/h*^*2*^ = 1, meaning that the proportion of phenotypic variance that is due to additive genetic effects is the same within- and between-populations. We note that this is a conservative estimation of *c/h*^*2*^ as many studies to-date assume *c*=1 and *h*^*2*^ =0.5 (e.g., Ghalambor et al. 2015). In our data, the 95% lower confidence limit of P_ST_ exceeds average F_ST_ at *c/h*^*2*^ > 1 for both modules B3 (Fig. 5A) and B36 (Fig. 5B). This result is robust to extremely conservative estimates of *c/h*^*2*^ <1 (Fig. 5A-B). Moreover, for most values of *c/h*^*2*^, P_ST_ exceeds even the upper 95% quantile of F_ST_ (Fig. 5A-B). By contrast, 95% confidence intervals for P_ST_ calculated on the B0 module (which is not associated with thermogenesis) overlap with F_ST_ at all values of *c/h*^*2*^ (Fig. 5C), indicating that differentiation in expression of this module does not exceed neutral expectations, as expected. P_ST_ calculated on VO_2_ across p0-p10 did not exceed F_ST_, except when we restricted it to p10 (Supplementary Fig. 2), the time point under which VO_2_ is significantly lower in highlander compared to lowlanders (Fig. 1A).

**Figure 5.**
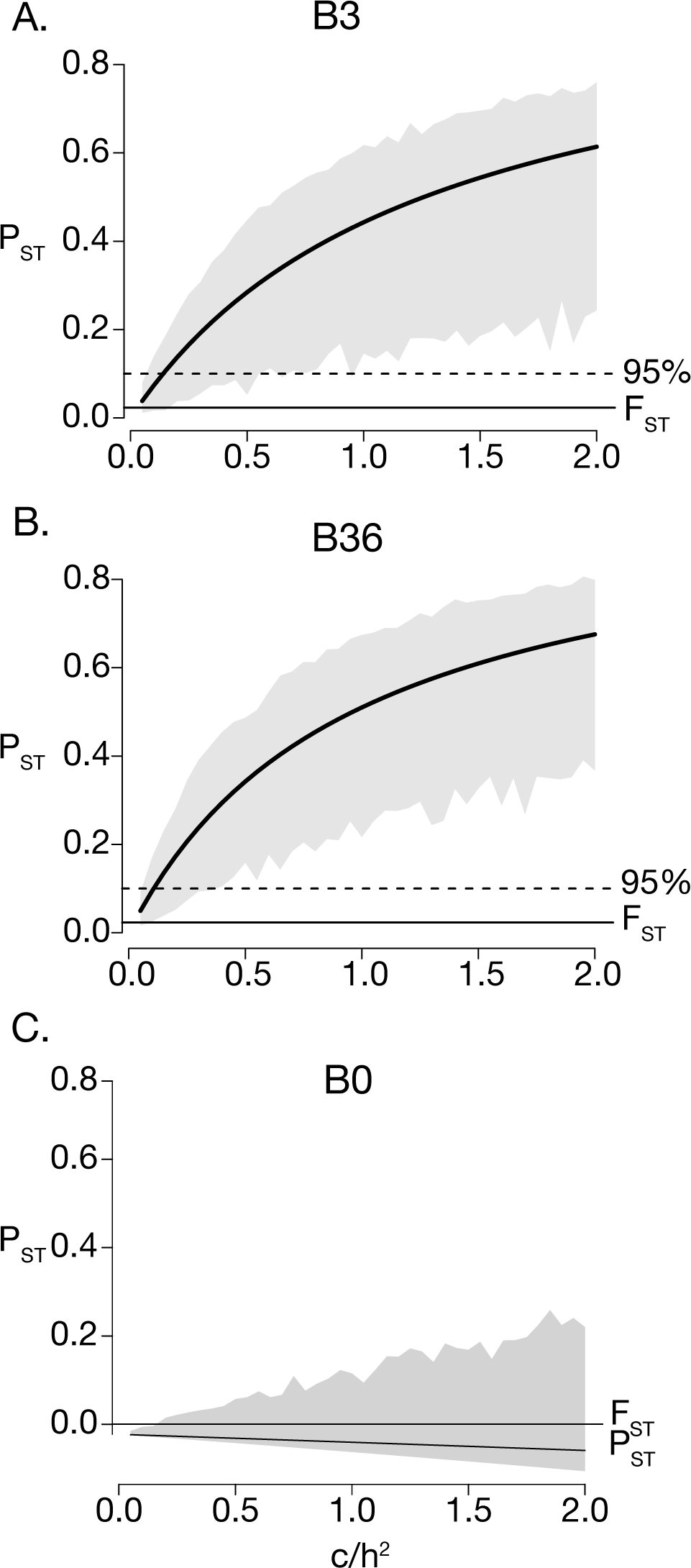
P_ST_ on module expression values for brown adipose tissue modules B0 (A), B3 (B), and B36 (C), which constitutes a suite of uncorrelated genes. P_ST_ values were calculated across a range of c/h^2^ values from 0-2. Grey polygons represent 95% confidence limits on P_ST_ generated from bootstrapping. Average F_ST_ and the upper 95% quantile of F_ST_ are shown. For nearly every value of P_ST_ for modules B3 and B36, but not B0, the lower confidence limit of P_ST_ of module expression is non-overlapping with F_ST_.

We found that P_ST_ on the skeletal muscle module M6 did not exceed neutral expectations: although observed P_ST_ does exceed F_ST_, lower 95% confidence limits overlap entirely with zero (Supplemental Fig. S1). We note that P_ST_ does not tend to overlap with the upper 95% limit of F_ST_ when we restrict it to p14 (the time point of significant differentiation in module expression and thermogenic capacity; Figs. 1B and 4B; see Supplemental Fig. S2). This pattern of phenotypic differentiation is mirrored by P_ST_ on thermogenic capacity at p14, which greatly exceeds the upper 95% limit of FST at all values of *c/h*^*2*^ (Supplemental Fig. S2).

Finally, we tested whether these candidate VO_2_-associated transcriptional modules were enriched for genes exhibiting signatures of positive natural selection, as measured by PBS. Roughly 25% of the genes in modules B3, B36 and M6 contained SNPs with PBS values that exceeded neutral expectations (Table 3). However, only B36 exhibited a significant enrichment of outlier SNPs relative to the background rate (Fisher’s Exact Test, *p*=0.02; Table 3). This analysis demonstrates that, although all phenotype-associated modules exhibit some proportion of genes under selection, only B36 contains more than was expected by chance.

**Table 3.**
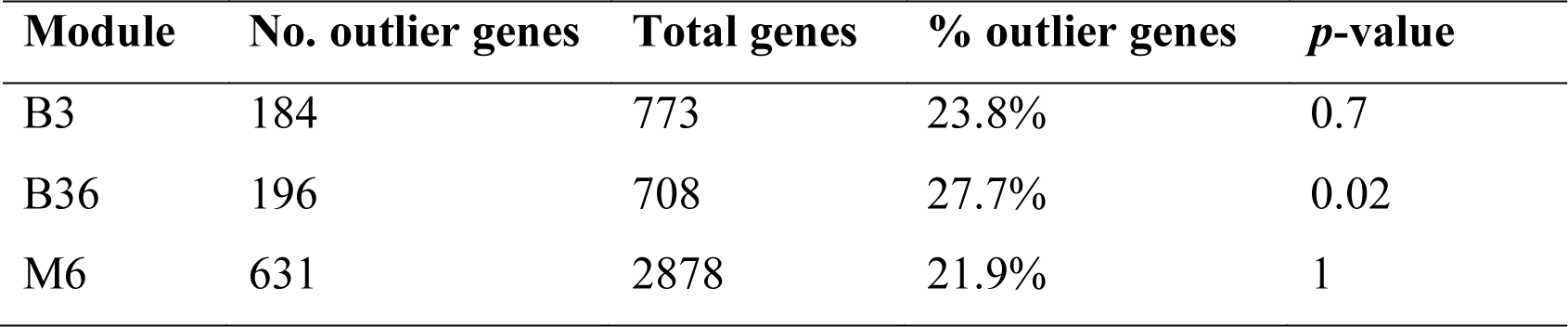
Number and proportion of genes in candidate modules containing outlier SNPs (PBS > 99.9% quantile of the simulated distribution). P-values obtained from Fisher’s Exact Tests

| **Module** | **No. outlier genes** | **Total genes** | **% outlier genes** | ***p*-value** |
| --- | --- | --- | --- | --- |
| B3 | 184 | 773 | 23.8% | 0.7 |
| B36 | 196 | 708 | 27.7% | 0.02 |
| M6 | 631 | 2878 | 21.9% | 1 |

## Discussion

We explored the mechanisms that underlie a developmental delay in the onset of thermogenesis in juvenile high-altitude deer mice (Fig 1). We show that the developmental trajectory of the thermoregulatory system has evolved at high altitude *via* selection on the regulatory systems that control the development of thermo-effector tissues, BAT and skeletal muscle. Combined with the results of Robertson et al. (2019) and Robertson and McClelland (2019), our work suggests that the cold hypoxic conditions of high-altitude force an alternative resource allocation strategy, whereby limited energy is put into developmental processes such as growth, over the development of the thermoregulatory machinery needed in heat production.

### A developmental delay in non-shivering thermogenesis is associated with a delay in expression of fuel supply, oxygenation, and axon guidance pathways in BAT

During early postnatal development mice rely exclusively on BAT for thermogenesis, and low-altitude pups activate BAT to maintain body temperature in the face of cold by p10. Our analysis suggests that an ontogenetic delay in the development of non-shivering thermogenesis in high-altitude deer mice (Fig. 1A) is attributable to a delay in the nervous system innervation and O_2_/fuel supply lines to a maturing BAT. WGCNA of BAT revealed two transcriptional modules (B3, B36) that are expressed at higher levels in lowlanders across age, but which show overall higher expression in both populations (Fig 3). Both modules were significantly correlated with cold-induced VO_2_, suggesting that changes in the expression of genes in these modules influence the development of thermogenesis. Module B3 was significantly enriched for pathways involved in metabolism, particularly fatty acid degradation (Fig. 3; Table S4). This is interesting since fat is the primary fuel source that powers uncoupling of cellular respiration from ATP production in non-shivering heat production (Cannon and Nedergaard 2004). Indeed, the β3 signaling cascade, initiated by cold, stimulates the lipolysis of fats used in β–oxidation and the eventual uncoupling of ATP synthesis from electron transport *via* activation of Uncoupling Protein 1 (*Ucp1*). Although not assigned to B3, we found that expression of *Ucp1* itself was lower in highlanders compared to lowlanders (ANOVA, main effect of population; F_1,30_=8.2; *p*=0.007).

Module B36, by contrast, was enriched for two pathways that are important in O_2_/fuel supply and cold-activation of BAT. For example, enrichment of the vasculature endothelial growth factor (VEGF) signaling pathway, and downregulation of *Vegfa* in highlanders (Fig 3B), suggests a reduction in the vascularization of BAT early in the postnatal period in high-altitude natives. This is notable since the VEGF cascade stimulates angiogenesis, increasing O_2_ and metabolic fuel delivery. Moreover, *Vegfa* itself contains two SNPs above the 99.9% PBS threshold (Schweizer, Velotta, et al. 2019), suggesting that changes in the VEGF pathway are the target of selection at high-altitude. We also detected enrichment of the KEGG pathway ‘axon guidance,’ which represents a key stage in development in which neurons send out axons to reach their correct targets. Indeed, BAT is activated by the sympathetic nervous system in response to cold, which leads to the release of norepinephrine and the signaling cascade that produces heat (Cannon and Nedergaard 2004).

A reduction in the expression of genes that participate in axon guidance suggests a developmental delay in the sympathetic innervation of BAT, which would inhibit recruitment and activation of this tissue during cold exposure (Robertson et al. 2019). Robertson et al. (2019) found lower levels of the enzyme tyrosine hydroxylase (TH) in the BAT of high-altitude pups at p10, lending support to the hypothesis that neurotransmitter synthesis and neural activation of BAT tissue is delayed in highlanders; TH is the rate-limiting enzyme in norepinephrine production and is present in sympathetic neurons innervating BAT. Related to the reduced sympathetic activation of BAT, we found that expression of the gene *calsyntenin 3* (*Clstn3*) is also downregulated in highlanders (ANOVA, main effect of population; F_1,30_=15.1; *p*<0.001). *Clstn3* is a known promoter of synapse formation; a recent study in house mice found that a mammalian-specific form (*Clstn3*β) enhances functional sympathetic innervation of BAT (Zeng et al. 2019).

Despite downregulation of genes involved in O_2_/fuel supply and innervation of BAT, Robertson at el. [2019] found that BAT mass does not differ between highlanders and lowlanders, and that citrate synthase (a biomarker of mitochondrial abundance) and UCP protein expression are likewise equivalent. Our results therefore suggest that, although BAT growth and metabolic potential may be equal in highlanders and lowlanders, a delay in the development of the innervation and O_2_/fuel supply to BAT may lead an inability of highlanders to respond to cold temperatures and thus mount an appropriate thermoregulatory response.

### A developmental delay in thermogenic capacity is associated with downregulation of energy metabolism pathways in muscle

Transcriptome analysis of gastrocnemius muscle revealed a large transcriptional module (M6) that closely tracked variation in thermogenic capacity (Fig. 4). These results suggest that the genes expressed in module M6 may underlie the developmental delay in thermogenesis beyond p10 (Fig. 1B). The M6 module (Table 2) was enriched for a host of functions related to skeletal muscle metabolism that reflect changes in its maturation with respect to shivering. Our analysis suggests that high-altitude mice show a developmental delay in the expression of nearly all pathways related to ATP generation by mitochondria, including glycolysis, TCA cycle, and oxidative phosphorylation (Fig 4C). Many of the genes that contribute to functional enrichment of metabolic pathways were related to the production of mitochondrial respiratory complex I, including 31 of 35 *Nduf* genes, suggesting that downregulation of metabolic pathways may be related to a delay in mitochondrial biogenesis. Indeed, the Gene Ontology terms “mitochondria and mitochondrial part” were also significantly enriched (Table S3).

At least two transcription factors that are key regulators of cellular metabolism were found in module M6 and were expressed at lower levels in highlanders (Fig. 4D-E). Estrogen-related receptor alpha, *Esrra*, for example, directs the expression of genes involved in mitochondrial biogenesis (Wu et al. 1999) and mitochondrial energy-producing pathways in skeletal muscle (Huss et al. 2004). Genes in module M6 were also enriched for *Esrra* binding sites (Table S3). Lowered expression of peroxisome proliferator activated receptor alpha (*Ppara*) suggests that fatty acid oxidation may also be depressed in highlanders, since this gene is a major regulator of fat metabolism capacity; functional terms related to fat oxidation however were not enriched in this module (Supplemental Table S4). Indeed, (Robertson and McClelland 2019) found that the ß-oxidation enzyme ß-hydroxyacyl-CoA dehydrogenase did not track changes in muscle aerobic capacity of highlanders over these ages. These data provide evidence that the developmental delay in thermogenic capacity is attributable to a delay in the expression of pathways that facilitate mitochondrial development and muscle aerobic capacity and allow for shivering thermogenesis.

### Evidence of natural selection

We found that population-level divergence in expression of VO_2_-associated transcriptional modules exceeded neutral expectations set by genetic differentiation (measured as genome-wide F_ST_), suggesting that population differences in expression may be adaptive. We expected that, if population-level divergence in module expression was adaptive, P_ST_ on module eigengene would exceed the upper bounds of the confidence intervals around F_ST_. At nearly every value of *c/h*^*2*^, the lower 95% limit of P_ST_ for BAT modules B3 and B36 exceeded average F_ST_ (Fig. 5). Moreover, at the null value of *c/h*^*2*^ =1 (Brommer 2011), P_ST_ was completely non-overlapping with the upper 95% quantile of F_ST_, indicating strong population-level differentiation (Fig. 5 A-B). This finding is consistent with P_ST_ on cold-induced VO_2_ values at p10 (Supplemental Fig. S2), the time point of greatest population differentiation (Fig. 1A). Although P_ST_ did not exceed F_ST_ for the skeletal muscle module M6 across all time points (p14-p27), we did find it to exceed neutral expectations when restricted to p14 (Supplemental Fig. S2). This is consistent with the finding that P_ST_ on thermogenic capacity also exceeds the upper 95% of F_ST_ at p14 only (Supplemental Fig. S2). These data together suggest that the delay in the onset of non-shivering and shivering thermogenesis, and their underlying regulatory control pathways, may be the target of selection at high-altitude.

We note that approximately 25% of genes in candidate modules contain at least on SNP above the PBS significance threshold (Table 3), and module B36 was significantly enriched for PBS outliers. Because PBS is polarized by the use of a second lowland population (see Methods), we can discern that these outlier loci are under positive selection at high-altitude specifically. Together these results support the interpretation that differentiation in candidate transcriptional modules, at least within BAT, is driven, in part, by natural selection.

### Summary and conclusions

We found that a developmental delay in the onset of independent thermoregulatory ability in deer mice native to high altitude is rooted in a delay in the expression of gene regulatory networks that contribute to sympathetic innervation of BAT that permits the response to cold, and in pathways that contribute to aerobic metabolism capabilities of both thermo-effector organs. Phenotypic divergence analysis suggests that both thermogenic delays and associated shifts in gene expression in BAT are driven by natural selection. An overabundance of genes exhibiting signatures of natural selection in BAT module B36 suggests that developmental changes to genes in this module may be particularly important in high-altitude adaptation. Our results demonstrate that changes to the developmental trajectory of thermoregulation at high-altitude are the result of regulatory delays in the development of thermo-effector organs, which limit the ability of high-altitude mice to respond to cold and mount a thermoregulatory response. Evidence presented here is consistent with the hypothesis that the developmental delay in thermogenesis at high altitude is attributable to an adaptive energetic trade-off (Robertson et al. 2019; Robertson and McClelland 2019). Although the exact cause is unknown, the combination of chronic cold and low O_2_ availability at high-altitude, coupled with differences in litter size that should affect competition among siblings for limited recourses, may contribute to the altered energetic conditions that drive this trade-off. Whatever the ultimate cause, our results suggest that high-altitude pups may benefit from allocating limited resources away from active thermoregulation, towards other developmental functions such as growth. Because high-altitude adult deer mice show the opposite pattern (*i.e.*, consistently higher thermogenic capacities, Cheviron et al. 2013), our combined results suggest that selection pressures at high altitude can have very different effects on the same physiological system depending on age. Future work on the selective drivers and energetic benefits of this putatively adaptive energy allocation strategy will shed further light on adaptive modification of developmental processes to meet the challenge of selection pressures that vary in strength and magnitude across ontogeny.

## Materials and Methods

### Animals and Experimental Procedures

Deer mice used in this study were the second-generation lab-reared descendants of two wild populations of *Peromyscus maniculatus* (Robertson et al. 2019; Robertson and McClelland 2019) raised under common garden conditions (24°C, 80 m a.s.l.) at McMaster University of Ontario, Canada. Individuals from the high-altitude population were trapped at the summit of Mount Evans in Clear Creek County CO, USA (4,350 m a.s.l.) and the low-altitude population from Nine-mile prairie, NE, USA (320 m a.s.l). Altricial rodents, such as deer mice, usually develop non-shivering thermogenesis (NST) prior to shivering due to the maturation of BAT which precedes the maturation of skeletal muscle. *P. maniculatus* are only capable of NST after the first 10 days postpartum (Robertson et al. 2019). In contrast, shivering develops between two and four weeks after birth (Robertson and McClelland 2019). We sampled BAT and skeletal muscle (specifically, the gastrocnemius) over these two periods. Briefly, pups were removed from the nest, then euthanized with an overdose of isoflurane followed by cervical dislocation. In pups ages 0, 2, 4, 6, 8 and 10 days postpartum the intrascapular depot of BAT was blunt dissected and cleaned of white adipose tissue. In pups aged 14, 21, and 27 days postpartum the gastrocnemius muscle of the lower hindlimb was blunt dissected. Tissues were flash frozen and stored at −80°C.

We re-analyzed cold-induced O_2_ consumption rate (VO_2_) data from Robertson et al. (2019) and Robertson and McClelland (2019) for low- and high-altitude native *P. maniculatus* only, omitting the low altitude congeneric, *P. leucopus*. Briefly, for pups aged 0-10 days postpartum cold-induced VO_2_ was assessed in response to a mild cold stress (10 minutes at 24°C). Pups were compared to a control litter mate who was maintained at 30°C for the same duration of trial in order to control for handling stress. For pups aged 14-27 days, maximum cold-induced VO_2_ (hereafter, thermogenic capacity) was measured using previously established methods for adult deer mice (Cheviron et al. 2012). VO_2_ max was induced by exposing pups to – 4°C in heliox air (21% O_2_ with He). All VO_2_ measurements were divided by mass to obtain mass-specific metabolic rates.

### Transcriptome analyses

We conducted high-throughput sequencing of RNA of BAT and gastrocnemius skeletal muscle in order to explore the mechanisms that underlie developmental delays in thermogenesis. Sample size varied from *n*=1-7 for each population, age, and tissue (see Table S1 in online supplementary material for full list). We assayed gene expression using TagSeq, a 3’ tag-based sequencing method following Lohman et al. (2016). First, we extracted RNA from <25 mg of tissue using TRI Reagent (Sigma-Aldrich), then assessed RNA quality using TapeStation (Agilent Technologies; RIN > 7). The Genome Sequencing and Analysis Facility at the University of Texas at Austin prepared TagSeq libraries, which were sequenced using Illumina HiSeq 2500. We filtered raw reads for length, quality, and PCR duplicates following Lohman et al. (2016) using scripts modified from those available online (https://github.com/z0on/tag-based_RNAseq). Using the FASTX-toolkit (http://hannonlab.cshl.edu/fastx_toolkit/), we cleaned and trimmed raw reads, which resulted in 3.2 million reads per individual. Filtered reads were then mapped to the *P. maniculatus* genome (NCBI GCA_000500345.1 Pman_1.0) using *bwa mem* (Li and Durbin 2010). We used *featureCounts* (Liao et al. 2014) to generate a table of transcript abundances. Since genes with low read counts are subject to measurement error (Robinson and Smyth 2007), we excluded those with less than an average of 10 reads per individual. We retained a total of 11,192 and 11,104 genes after filtering for BAT and skeletal muscle transcriptomes, respectively. Five outlier samples (one BAT, four skeletal muscle) were removed following visual inspection of multi-dimensional scaling plots using *plotMDS* in *edgeR* following Chen et al. (2019).

We assessed overall patterns of gene expression among BAT and skeletal muscle transcripts using Principal Components Analysis (PCA), while weighted gene co-expression network analysis (WGCNA v. 1.41-1; Langfelder and Horvath 2008) was used to identify potential regulatory mechanisms that underlie variation in thermogenesis across developmental time. PCA and WGCNA analyses were conducted separately on BAT and skeletal muscle samples in R v. 2.5.2 (R Core Team). WGCNA identified clusters of genes with highly correlated expression profiles (hereafter, modules). This approach was successfully implemented in recent studies that aimed to relate gene expression with higher-level phenotypes (Plachetzki et al. 2014; Velotta et al. 2016; Velotta et al. 2018). Prior to performing WGCNA, we normalized raw read counts by library size and log-transformed them using the functions *calcNormFactors* and *cpm*, respectively, from the R package *edgeR* (Robinson et al. 2010). Module detection was performed using the *blockwiseModules* function in WGCNA with networkType set to “signed” (Langfelder and Horvath 2008). Briefly, Pearson correlations of transcript abundance data were calculated between pairs of genes, after which an adjacency matrix was computed by raising the correlation matrix to a soft thresholding power of β = 6. Soft thresholding power β is chosen to achieve an approximately scale free topology, an approach that favors strong correlations (Zhang and Horvath 2005). We chose a β since it represents the value for which improvement of scale free topology model fit begins to decrease with increasing thresholding power. A topological overlap measure was computed from the resulting adjacency matrix for each gene pair. Topologically-based dissimilarity was then calculated and used as input for average linkage hierarchical clustering in the creation of cluster dendrograms for both left and right ventricles. Modules were identified as branches of the resulting cluster tree using the dynamic tree-cutting method (Langfelder and Horvath 2008). We assigned modules a unique identification according to tissue (B0-B37 for brown adipose tissue; M0-M7 for skeletal muscle). Genes that could not be clustered into a module were given the designation B0 and M0 for BAT and skeletal muscle, respectively.

Once modules were defined, we used a multi-step process to associate expression among individuals with variation in VO_2_. First, we summarized module expression using principal components analysis (PCA) of gene expression profiles (*blockwiseModules* in WGCNA); because genes within modules are highly correlated by definition, the first principal component (referred to as the module eigengene value) was used to represent module expression (Langfelder and Horvath 2008). We used module eigengene values to test for associations between module expression and VO_2_ for each module (Pearson correlation; *cor* function in WGCNA) in the BAT network. *P*-values for the correlation were determined by a Student’s asymptotic test (*corPvalueStuden*t in WGCNA). For skeletal muscle, association tests were conducted on population-age means since VO_2_ measurements do not exist for the individuals sequenced. Among phenotype-associated modules, we conducted analysis of variance (ANOVA) on rank-transformed module eigengene values in order to test for the effects of population, age, and their interaction on module expression. P-values from association tests and ANOVAs were corrected for multiple testing using the false-discovery rate method.

We performed functional enrichment analysis on all modules exhibiting either a statistical association with VO_2_, or a significant effect in ANOVA, or both, using the R package *gProfilerR* (Reimand et al. 2016). *Peromyscus maniculatus* gene names were converted to *Mus musculus* gene names using a custom script (Online supplementary material). We corrected for multiple testing using *gProfilerR’s* native g:SCS algorithm. We used the list of genes from filtered BAT and gastroc transcriptomes as a custom background list in all enrichment analyses. We searched for enrichment among terms from Gene Ontology (GO; The Gene Ontology Consortium 2019), KEGG (Kanehisa and Goto 2000), Reactome (Fabregat et al. 2018), and WikiPathways (Slenter et al. 2018) databases, as well as transcription factor binding sites from the TRANSFAC database (Matys et al. 2006).

### Tests of phenotypic and genetic divergence

We used two complementary approaches to assess phenotypic and genetic divergence to test for signatures of natural selection in VO_2_-associated modules. To assess phenotypic divergence, we calculated P_ST_ on module eigengene values following Brommer (2011) using Eq. (1),

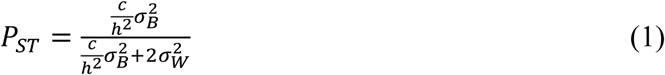

where 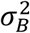 denotes between-population phenotypic variance, 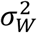 denotes within-population phenotypic variance, *h*^*2*^ the narrow-sense heritability (the proportion of phenotypic variance that is due to additive genetic effects), and *c* a scalar representing the total phenotypic variance due to additive genetic effects across populations. Within- and between-population variances were estimated from ANOVA, where module eigengene value were response variables, and postnatal day, population, and family nested within population were included as main effects. Within-population variance 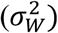 was calculated as the residual mean squares (MS) from the ANOVA model. Between-population variance was estimated after Antoniazza et al (2010) following Eq. (2),

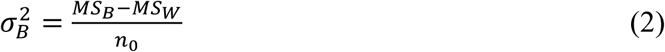

where *MS*_*B*_ and *MS*_*W*_ are the population effect and the residual mean square variances from the ANOVA model, respectively, and *n*_0_ a weighted average of sample size based on Sokal and Rohlf (1995) following Eq. (3).

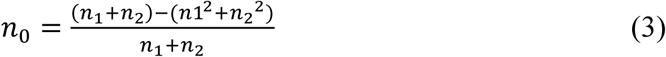

where *n*_*1*_ and *n*_*2*_ are the sample sizes of population one and two, respectively. The ratio c/h^2^ was unknown and could not be readily estimated using our experimental design. As such, we calculated P_ST_ across a range of values of c/h^2^ from 0 (no heritability) to 2, which is where the relationship between c/h^2^ and P_ST_ begins to reach asymptote (Brommer 2011; Fig. 5). We calculated 95% confidence intervals for each value of P_ST_ calculated across values of c/h^2^ ranging from 0-2 at 0.05 level increments (total of 40 P_ST_ estimates).

P_ST_ estimates were compared to neutral divergence expectations set by between-population differentiation. We calculated pairwise genetic differentiation between the lowland and highland populations using F_ST_ (estimated with Weir’s Theta; Weir and Cockerham 1984) and previously sequenced exome data (Schweizer, Velotta, et al. 2019). Briefly, F_ST_ was calculated for approximately 5 million high-quality bi-allelic single nucleotide polymorphisms (SNPs) using the ‘--weir-fst-pop’ flag within vcftools v0.1.16 (Danecek et al. 2011). We compared P_ST_ of VO_2_-associated modules to that of “Module 0,” which contains the suite of genes that could not be reliably clustered (B0 in BAT; M0 in skeletal muscle). The null expectation is that P_ST_ should not exceed the neutral expectation set by F_ST_ at any value of c/h^2^. We expected that, if natural selection has shaped shifts in gene expression, then P_ST_ would exceed F_ST_ in VO_2_-associated modules, but not in Modules B0 or M0 (which serve as a measure of background levels of expression differentiation). We calculated P_ST_ on cold-induced VO_2_ (p0-p10) and thermogenic capacity (p14-p27) as above in order to determine whether these higher level phenotypic traits were also the targets of natural selection.

Finally, we took a population genomic approach to assess whether genes in VO_2_-associated transcriptional modules exhibit signatures of positive natural selection at high-altitude. To do this, we re-analyzed Population Branch Statistic (PBS; Yi et al. 2010) data from (Schweizer, Velotta, et al. 2019). Here, PBS identifies loci that exhibit extreme allele frequency differences in highland (Mt. Evans) relative to lowland (Lincoln and Merced, CA) populations. We previously calculated PBS for approximately 5 million exome-wide bi-allelic SNPs, then calculated the demographically-corrected significance of PBS values using a simulated distribution of 500,000 SNPs (see Schweizer, Velotta, et al. 2019 for simulation details). We first determined significant outlier SNPs with a PBS value located above the 99.9% quantile, then identified those SNPs located within genes belonging to VO_2_-associated modules using Ensembl’s Variant Effect Predictor (McLaren et al. 2010) with the *P. maniculatus* reference genome annotation data set. We used Fisher’s Exact Tests to determine whether thermogenesis-associated modules were statistically enriched for loci that exceeded the 99.9% threshold.

## Acknowledgements

We thank Maria Stager and Timothy Moore for help with statistical analyses. Thank you to Kamilla Bentsen and Madilyn Head for help extracting RNA. Funding to JPV provided by the National Institutes of Health, National Heart, Lung, and Blood Institute, Research Service Award Fellowship (1F32HL136124-01). Funding to RMS and ZAC provided by the National Science Foundation (Postdoctoral Research Fellowship in Biology 1612859 to RMS, IOS-1755411 and OIA 1736249 to ZAC). Funding to GBM provided by the National Sciences and Engineering Research Council of Canada (RGPIN 462246-2014).

## Supplementary Material

**Table S1.** Sample sizes for tag-seq analysis after outlier removal (see Methods).

|  | p0 | p2 | p6 | p8 | p10 | p14 | p21 | p27 |
| --- | --- | --- | --- | --- | --- | --- | --- | --- |
| Lowland | 2 | 4 | 4 | 4 | 6 | 3 | 3 | 5 |
| Highland | 1 | 2 | 2 | 3 | 6 | 4 | 5 | 6 |

**Table S2.** Table of all genes expressed in brown adipose tissue (BAT) and skeletal muscle and their module assignments using WGCNA. Modules were given color name identifiers by WGCNA, which were converted into numbered identifiers (B# for brown adipose tissue; M# for skeletal muscle). Gene names derived from the *Peromyscus maniculatus* genome Annotation Release 100.

**Table S3.** ANOVA and association analysis results for BAT modules. Table presents F-values and false-discovery rate corrected *p*-values for main effects of population, postnatal age, and their interaction from the ANOVA on rank-transformed module expression values. Pearson correlation coefficients and false-discovery rate corrected *p*-values for the association between VO_2_ and module expression are also presented.

**Table S4.** Results of *gProfileR* functional enrichment analysis of phenotype-associated BAT and skeletal muscle WGCNA modules. Source represents whether terms were identified in the Gene Ontology (GO) database (BP: Biological Process, CC: Cellular Component, MF: Molecular Function), Kyoto Encyclopedia of Genes and Genomes (KEGG) pathways, Reactome pathways (REAC), Wikipathways (WP), or transcription factor binding sites identified in the TRANSFAC TFBS database (TF). ‘Adjusted_p_value’ is the corrected *p*-value for the enrichment analysis – we corrected for multiple testing using *gProfilerR’s* native g:SCS algorithm. ‘Term_size’ represents the number of genes in each functional category term, while ‘query_size’ represents the number of searchable genes in each module. The column ‘intersections’ lists the gene query names associated with each term. Modules without reported terms had none significantly enriched at *p*> 0.05 after correction for multiple testing.

**Figure S1.**
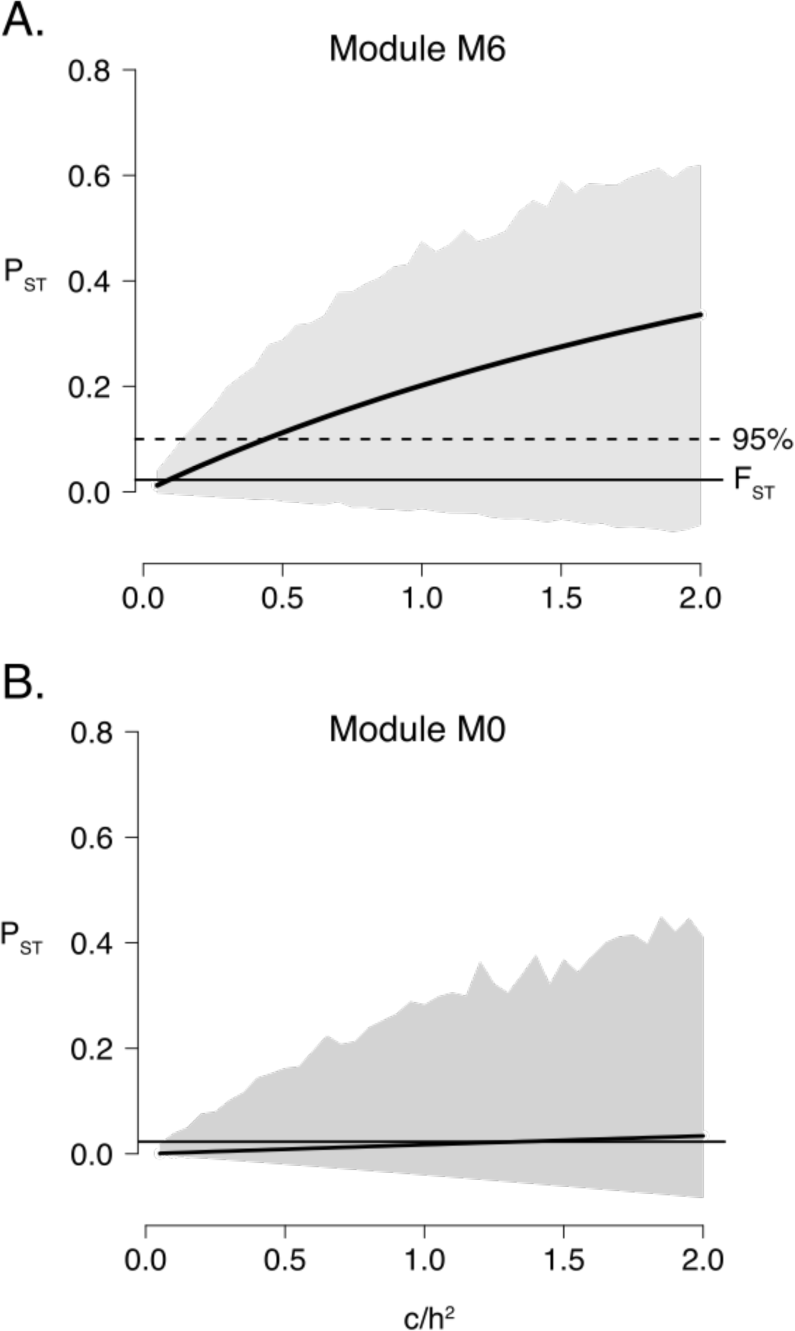
P_ST_ on module expression values for skeletal muscles modules M6 (A), and M0, which constitutes the suite of uncorrelated genes. P_ST_ values were calculated across a range of c/h^2^ values from 0-2. Grey polygons represent 95% confidence limits on P_ST_ generated from bootstrapping. Average F_ST_ and the upper 95% quantile of F_ST_ are shown.

**Figure S2.**
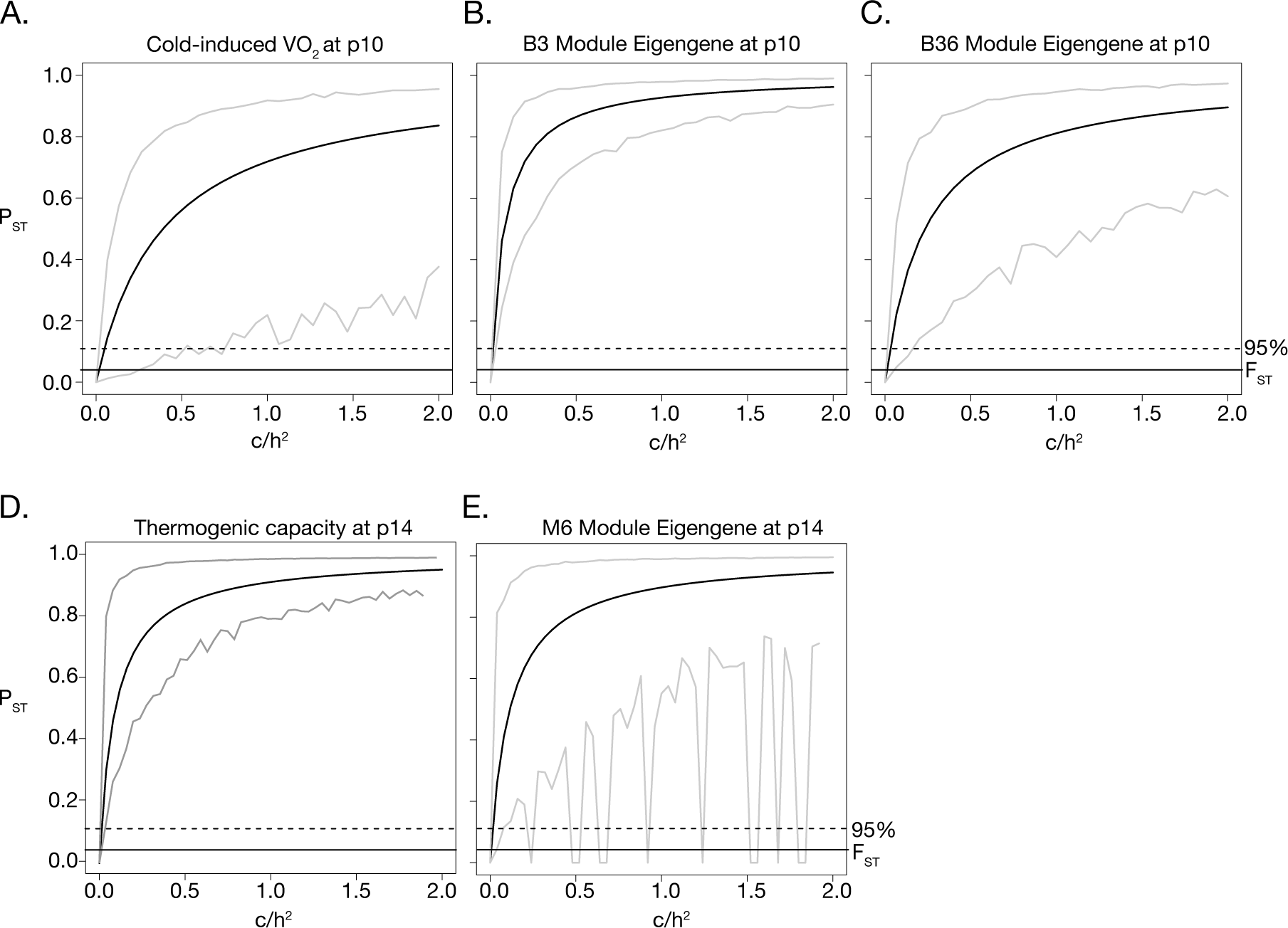
P_ST_ on cold-induced VO_2_ (A) and thermogenic capacity data (D) at the time point of greatest population differentiation (p10 and p14, respectively). Also plotted is PST on candidate module eigengene values B3 (B), B36 (C), and M6 (E) for at the same time points. P_ST_ values were calculated across a range of *c/h*^*2*^ values from 0-2. Grey polygons represent 95% confidence limits on P_ST_ generated from bootstrapping. Average F_ST_ and the upper 95% quantile of F_ST_ are shown.

## References

Antoniazza S, Burri R, Fumagalli L, Goudet J, Roulin A. 2010. Local adaptation maintains clinal vriation in melanin-based coloration of European barn owls (*Tyto alba*). Evolution 64:1944–1954.

Arjamaa O, Lagerspetz KYH. 1979. Postnatal development of shivering in the mouse. Journal of Thermal Biology 4:35–39.

Beall CM. 2007. Two routes to functional adaptation: Tibetan and Andean high-altitude natives. PNAS 104:8655–8660.

Bouverot P. 1985. Adaptation to Altitude-Hypoxia in Vertebrates. Springer Science & Business Media

Brommer JE. 2011. Whither Pst? The approximation of Qst by Pst in evolutionary and conservation biology. Available from: https://onlinelibrary.wiley.com/doi/full/10.1111/j.1420-9101.2011.02268.x

Campbell-Staton SC, Bare A, Losos JB, Edwards SV, Cheviron ZA. 2018. Physiological and regulatory underpinnings of geographic variation in reptilian cold tolerance across a latitudinal cline. Molecular Ecology 27:2243–2255.

Campbell-Staton SC, Cheviron ZA, Rochette N, Catchen J, Losos JB, Edwards SV. 2017. Winter storms drive rapid phenotypic, regulatory, and genomic shifts in the green anole lizard. Science 357:495–498.

Cannon B, Nedergaard J. 2004. Brown adipose tissue: function and physiological significance. Physiol. Rev. 84:277–359.

Chappell MA, Snyder LR. 1984. Biochemical and physiological correlates of deer mouse alpha-chain hemoglobin polymorphisms. PNAS 81:5484–5488.

Chen Y, McCarthy DJ, Ritchie M, Robinson M, Smyth GK. 2019. edgeR: differential expression analysis of digital gene expression data User’s Guide. Available from: http://scholar.googleusercontent.com/scholar?q=cache:wKdKwEukm7wJ:scholar.google.com/+edgeR+and+outlier&hl=en&as_sdt=0,27

Cheviron ZA, Bachman GC, Connaty AD, McClelland GB, Storz JF. 2012. Regulatory changes contribute to the adaptive enhancement of thermogenic capacity in high-altitude deer mice. PNAS 109:8635–8640.

Cheviron ZA, Bachman GC, Storz JF. 2013. Contributions of phenotypic plasticity to differences in thermogenic performance between highland and lowland deer mice. J. Exp. Biol. 216:1160–1166.

Cheviron ZA, Connaty AD, McClelland GB, Storz JF. 2014. Functional genomics of adaptation to hypoxic cold-stress in high-altitude deer mice: transcriptomic plasticity and thermogenic performance. Evolution 68:48–62.

Danecek P, Auton A, Abecasis G, Albers CA, Banks E, DePristo MA, Handsaker RE, Lunter G, Marth GT, Sherry ST, et al. 2011. The variant call format and VCFtools. Bioinformatics 27:2156–2158.

DeBiasse MB, Kelly MW. 2016. Plastic and Evolved Responses to Global Change: What Can We Learn from Comparative Transcriptomics? J Hered 107:71–81.

Fabregat A, Jupe S, Matthews L, Sidiropoulos K, Gillespie M, Garapati P, Haw R, Jassal B, Korninger F, May B, et al. 2018. The Reactome Pathway Knowledgebase. Nucleic Acids Res. 46:D649–D655.

Garland T, Carter PA. 1994. Evolutionary Physiology. Annual Review of Physiology 56:579–621.

Garland T, Losos JB. 1994. Ecological morphology of locomotor performance in squamate reptiles. In: Ecological morphology: integrated organismal biology (Wainwright PC, Reilly SM, eds). Chicago, IL: Chicago University Press. p. 240–302.

Ghalambor CK, Hoke KL, Ruell EW, Fischer EK, Reznick DN, Hughes KA. 2015. Non-adaptive plasticity potentiates rapid adaptive evolution of gene expression in nature. Nature 525:372–375.

Hayes JP, O’Connor CS. 1999. Natural selection on thermogenic capacity of high-altitude deer mice. Evolution 53:1280–1287.

Hill RW. 1983. Thermal Physiology and Energetics of Peromyscus; Ontogeny, Body Temperature, Metabolism, Insulation, and Microclimatology. J Mammal 64:19–37.

Huey RB, Stevenson RD. 1979. Integrating thermal physiology and ecology of cctotherms: A discussion of approaches. Integr Comp Biol 19:357–366.

Huss JM, Torra IP, Staels B, Giguère V, Kelly DP. 2004. Estrogen-related receptor α directs peroxisome proliferator-activated receptor α signaling in the transcriptional control of energy metabolism in cardiac and skeletal muscle. Mol Cell Biol 24:9079–9091.

Irschick DJ, Garland T. 2001. Integrating function and ecology in studies of adaptation: Investigations of locomotor capacity as a model system. Annual Review of Ecology and Systematics 32:367–396.

Irschick DJ, Meyers JJ, Husak JF, Galliard J-FL. 2008. How does selection operate on whole-organism functional performance capacities? A review and synthesis. Evol. Ecol. Res. 10:177–197.

Issemann I, Green S. 1990. Activation of a member of the steroid hormone receptor superfamily by peroxisome proliferators. Nature 347:645–650.

Ivy CM, Scott GR. 2017. Control of breathing and ventilatory acclimatization to hypoxia in deer mice native to high altitudes. Acta Physiol 221:266–282.

Kanehisa M, Goto S. 2000. KEGG: Kyoto Encyclopedia of Genes and Genomes. Nucleic Acids Res 28:27–30.

King JA. 1968. Biology of Peromyscus (Rodentia). Biology of Peromyscus (Rodentia). [Internet]. Available from: https://www.cabdirect.org/cabdirect/abstract/19710101319

Lagerspetz KYH. 1966. Postnatal development of thermoregulation in laboratory mice. Helgoländer wissenschaftliche Meeresuntersuchungen 14.

Lailvaux SP, Husak JF. 2014. The life history of whole-organism performance. The Quarterly Review of Biology 89:285–318.

Langfelder P, Horvath S. 2008. WGCNA: an R package for weighted correlation network analysis. BMC Bioinformatics 9:559.

Lau DS, Connaty AD, Mahalingam S, Wall N, Cheviron ZA, Storz JF, Scott GR, McClelland GB. 2017. Acclimation to hypoxia increases carbohydrate use during exercise in high-altitude deer mice. American Journal of Physiology-Regulatory, Integrative and Comparative Physiology 312:R400–R411.

Li H, Durbin R. 2010. Fast and accurate long-read alignment with Burrows-Wheeler transform. Bioinformatics 26:589–595.

Liao Y, Smyth GK, Shi W. 2014. featureCounts: an efficient general purpose program for assigning sequence reads to genomic features. Bioinformatics 30:923–930.

Lohman BK, Weber JN, Bolnick DI. 2016. Evaluation of TagSeq, a reliable low-cost alternative for RNAseq. Mol Ecol Resour:n/a-n/a.

Lui MA, Mahalingam S, Patel P, Connaty AD, Ivy CM, Cheviron ZA, Storz JF, McClelland GB, Scott GR. 2015. High-altitude ancestry and hypoxia acclimation have distinct effects on exercise capacity and muscle phenotype in deer mice. Am. J. Physiol. Regul. Integr. Comp. Physiol. 308:R779–R791.

Mahalingam S, McClelland GB, Scott GR. 2017. Evolved changes in the intracellular distribution and physiology of muscle mitochondria in high-altitude native deer mice. The Journal of Physiology 595:4785–4801.

Matys V, Kel-Margoulis OV, Fricke E, Liebich I, Land S, Barre-Dirrie A, Reuter I, Chekmenev D, Krull M, Hornischer K, et al. 2006. TRANSFAC® and its module TRANSCompel®: transcriptional gene regulation in eukaryotes. Nucleic Acids Res 34:D108–D110.

McLaren W, Pritchard B, Rios D, Chen Y, Flicek P, Cunningham F. 2010. Deriving the consequences of genomic variants with the Ensembl API and SNP Effect Predictor. Bioinformatics 26:2069–2070.

Nikel KE, Shanishchara NK, Ivy CM, Dawson NJ, Scott GR. 2018. Effects of hypoxia at different life stages on locomotory muscle phenotype in deer mice native to high altitudes. Comparative Biochemistry and Physiology Part B: Biochemistry and Molecular Biology 224:98–104.

Pembrey MS. 1895. The effect of variations in external temperature upon the output of carbonic acid and the temperature of young animals. J Physiol 18:363–379.

Plachetzki DC, Pankey MS, Johnson BR, Ronne EJ, Kopp A, Grosberg RK. 2014. Gene co-expression modules underlying polymorphic and monomorphic zooids in the colonial hydrozoan, *Hydractinia symbiolongicarpus*. Integr. Comp. Biol.:icu080.

Reimand J, Arak T, Adler P, Kolberg L, Reisberg S, Peterson H, Vilo J. 2016. g:Profiler—a web server for functional interpretation of gene lists (2016 update). Nucl. Acids Res. 44:W83–W89.

Robertson CE, McClelland GB. 2019. Postnatal maturation of skeletal muscle drives adaptive thermogenic capacity of high-altitude deer mice, *Peromyscus maniculatus*. Journal of Experimental Biology.

Robertson CE, Tattersall GJ, McClelland GB. 2019. Development of homeothermic endothermy is delayed in high-altitude native deer mice (Peromyscus maniculatus). Proceedings of the Royal Society B: Biological Sciences 286:20190841.

Robinson MD, McCarthy DJ, Smyth GK. 2010. edgeR: a Bioconductor package for differential expression analysis of digital gene expression data. Bioinformatics 26:139–140.

Robinson MD, Smyth GK. 2007. Moderated statistical tests for assessing differences in tag abundance. Bioinformatics 23:2881–2887.

Schweizer RM, Velotta JP, Ivy CM, Jones MR, Muir SM, Bradburd GS, Storz JF, Scott GR, Cheviron ZA. 2019. Physiological and genomic evidence that selection on the transcription factor *Epas1* has altered cardiovascular function in high-altitude deer mice. PLOS Genetics 15:e1008420–e1008420.

Scott GR, Elogio TS, Lui MA, Storz JF, Cheviron ZA. 2015. Adaptive modifications of muscle phenotype in high-altitude deer mice are associated with evolved changes in gene regulation. Mol. Biol. Evol. 32:1962–1976.

Simonson TS, McClain DA, Jorde LB, Prchal JT. 2011. Genetic determinants of Tibetan high-altitude adaptation. Hum Genet 131:527–533.

Slenter DN, Kutmon M, Hanspers K, Riutta A, Windsor J, Nunes N, Mélius J, Cirillo E, Coort SL, Digles D, et al. 2018. WikiPathways: a multifaceted pathway database bridging metabolomics to other omics research. Nucleic Acids Res 46:D661–D667.

Snyder LRG. 1981. Deer mouse hemoglobins: is there genetic adaptation to high altitude? BioScience 31:299–304.

Snyder LRG, Born S, Lechner AJ. 1982. Blood oxygen affinity in high- and low-altitude populations of the deer mouse. Resp. Physiol. 48:89–105.

Sokal RR, Rohlf FJ. 1995. Biometry. 3rd ed. New York, NY: W.H.H.H Freeman and Company

Storz JF. 2007. Hemoglobin Function and Physiological Adaptation to Hypoxia in High-Altitude Mammals. J. Mammal. 88:24–31.

Storz JF. 2016. Hemoglobin–oxygen affinity in high-altitude vertebrates: is there evidence for an adaptive trend? Journal of Experimental Biology 219:3190–3203.

Storz JF, Scott GR, Cheviron ZA. 2010. Phenotypic plasticity and genetic adaptation to high-altitude hypoxia in vertebrates. J Exp Biol 213:4125–4136.

Tate KB, Ivy CM, Velotta JP, Storz JF, McClelland GB, Cheviron ZA, Scott GR. 2017. Circulatory mechanisms underlying adaptive increases in thermogenic capacity in high-altitude deer mice. Journal of Experimental Biology 220:3616–3620.

Velotta JP, Jones J, Wolf CJ, Cheviron ZA. 2016. Transcriptomic plasticity in brown adipose tissue contributes to an enhanced capacity for nonshivering thermogenesis in deer mice. Mol Ecol 25:2870–2886.

Velotta JP, McCormick SD, Jones AW, Schultz ET. 2018. Reduced Swimming Performance Repeatedly Evolves on Loss of Migration in Landlocked Populations of Alewife. Physiological and Biochemical Zoology 91:814–825.

Velotta JP, Wegrzyn JL, Ginzburg S, Kang L, Czesny S, O’Neill RJ, McCormick SD, Michalak P, Schultz ET. 2017. Transcriptomic imprints of adaptation to fresh water: parallel evolution of osmoregulatory gene expression in the Alewife. Mol Ecol 26:831–848.

Weir BS, Cockerham CC. 1984. Estimating F-Statistics for the Analysis of Population Structure. Evolution 38:1358–1370.

Whitehead A. 2012. Comparative genomics in ecological physiology: toward a more nuanced understanding of acclimation and adaptation. J. Exp. Biol. 215:884–891.

Wu Z, Puigserver P, Andersson U, Zhang C, Adelmant G, Mootha V, Troy A, Cinti S, Lowell B, Scarpulla RC, et al. 1999. Mechanisms controlling mitochondrial biogenesis and respiration through the thermogenic coactivator PGC-1. Cell 98:115–124.

Yi X, Liang Y, Huerta-Sanchez E, Jin X, Cuo ZXP, Pool JE, Xu X, Jiang H, Vinckenbosch N, Korneliussen TS, et al. 2010. Sequencing of 50 Human Exomes Reveals Adaptation to High Altitude. Science 329:75–78.

Zeng X, Ye M, Resch JM, Jedrychowski MP, Hu B, Lowell BB, Ginty DD, Spiegelman BM. 2019. Innervation of thermogenic adipose tissue via a calsyntenin 3β–S100b axis. Nature 569:229.

Zhang B, Horvath S. 2005. A general framework for weighted gene co-expression network analysis. Statistical Applications in Genetics and Molecular Biology [Internet] 4. Available from: http://www.degruyter.com/view/j/sagmb.2005.4.1/sagmb.2005.4.1.1128/sagmb.2005.4.1.1128.xml

